# Perivascular Secretome Influences Hematopoietic Stem Cell Maintenance in a Gelatin Hydrogel

**DOI:** 10.1101/2020.04.25.061572

**Authors:** Victoria Barnhouse, Nathan Petrikas, Cody Crosby, Janet Zoldan, Brendan Harley

## Abstract

Adult hematopoietic stem cells (HSCs) produce the body’s full complement of blood and immune cells. They reside in specialized microenvironments, or niches, within the bone marrow. The perivascular niche near blood vessels is believed to help maintain primitive HSCs in an undifferentiated state but demonstration of this effect is difficult. *In vivo* studies make it challenging to determine the direct effect of the endosteal and perivascular niches as they can be in close proximity, and two-dimensional *in vitro* cultures often lack an instructive extracellular matrix environment. We describe a tissue engineering approach to develop and characterize a three-dimensional perivascular tissue model to investigate the influence of the perivascular secretome on HSC behavior. We generate 3D endothelial networks in methacrylamide-functionalized gelatin hydrogels using human umbilical vein endothelial cells (HUVECs) and mesenchymal stromal cells (MSCs). We identify a subset of secreted factors important for HSC function, and examine the response of primary murine HSCs in hydrogels to the perivascular secretome. Within 4 days of culture, perivascular conditioned media promoted maintenance of a greater fraction of hematopoietic stem and progenitor cells. This work represents an important first-generation perivascular model to investigate the role of niche secreted factors on the maintenance of primary HSCs.

## 1. Introduction

Adult hematopoietic stem cells (HSCs) reside in the bone marrow and are responsible for producing the body’s full complement of blood and immune cells through a process termed hematopoiesis.^6^ Clinically, HSCs are used to treat a number of hematological disorders including leukemia and lymphoma via transplantation of donor HSCs from either autologous, allogenic, or umbilical cord blood sources into the patient after successful ablation of the patient’s bone marrow.^13,18^ However there is a high rate of failure associated with unsuccessful initial engraftment, loss of donor cells following engraftment, or low stem cell dose.^56^ Increased success has been correlated with an increased number of donor HSCs infused, and thus methods of expansion prior to transplant has been an active area of investigation.^19^ Currently, there is no reliable method of expanding HSCs without leading to exhaustion following transplant. The differentiation hierarchy of HSCs is thought to consist of multiple stem cell types and early progenitors. A small, rare population of long-term repopulating HSCs (LT-HSCs) are largely quiescent, meaning they exit the cell cycle and divide only rarely, and have the capacity for continued self-renewal. Comparatively, short-term repopulating HSC (ST-HSC) possess more limited self-renewal capacity, while multipotent progenitor cells (MPPs) represent an early progenitor cell that retains the ability to differentiate into either the myeloid or lymphoid lineages but lack self-renewal capacity. Collectively, these three cell types are termed hematopoietic stem and progenitors (HSPCs) and can be distinguished as the Lin^−^ Sca-1^+^ c-kit^+^ (LSK) population via flow cytometry.^12,73^

The stem cell microenvironment, termed niche, provides signals to influence HSC fate decisions of quiescence, self-renewal, and differentiation. The bone marrow contains several distinct niche environments hypothesized to impart disparate hematopoietic functions, including the endosteal niche near the surface of bone and the perivascular niche, near blood vessels in the marrow. The perivascular niche (<20μm from vessels) has been suggested to be important for primitive HSCs, with *in vivo* imaging showing close association with vessels in the marrow.^1,15,42,53^ Further investigation has shown that the perivascular niche provides biomolecular signals that can influence HSC function (i.e., SCF, FGF2, IGFBP2, DLL1) implicating the perivascular secretome in influencing HSC fate decisions.^21,40,71^ Many such studies have been performed *in vivo*, where a complex microenvironment makes it difficult to observe the direct effects of any experimental conditions.^2,21,34,42^ Tissue engineering platforms provide the potential to control the complexity of the extracellular environment in order to parse out effects of multiple niche components. These tissue engineering approaches have been used to identify regulators of HSC function in culture and gained valuable insight into the role of various niche factors. *Poulos et al*. demonstrated that bone marrow endothelial cells can support functional HSCs *in vitro* using a 2D co-culture model.^60^ Recently, *Braham et al* investigated multiple combinations of stromal cells in a hydrogel system, and found that optimal support of hematopoietic stem and progenitor cells (HSPCs) was provided by adipogenic and osteogenic cells, along with endothelial cells without the need for supplemental cytokine addition.^5^

Our lab has previously developed a methacrylamide-functionalized gelatin (GelMA) hydrogel platform to probe questions related to biomolecule presentation and biotransport properties in an artificial HSC niche.^26,44,57^ The GelMA hydrogel provides RGD binding motifs and are sensitive to MMP-mediated degradation. These studies have shown that immobilized SCF better maintained primitive HSCs, and a co-culture of MSCs and HSCs in an initially low-diffusivity hydrogel promotes the maintenance of quiescent early hematopoietic cells.^26,44^ We have recently adapted this hydrogel to generate robust endothelial networks as a perivascular model for studies of glioblastoma within our hydrogel system, using human umbilical vein endothelial cells (HUVECs) and normal human lung fibroblasts (NHLFs).^48^ Therefore, the objective of this study was to analyze the biomolecular signals from a perivascular model and determine their effect on the maintenance and differentiation of primary murine HSCs. We report the ability to monitor the formation of stable microvascular networks *in vitro*, analyze the content and hematopoietic-relevance of the secretome generated by these endothelial cell networks, then examine the role of these factors on the maintenance of primary HSCs *in vitro*.

## 2. Materials and Methods

### 2.1. Cell Culture

HUVECs and hMSCs were purchased from Lonza (Walkersville, MD). HUVECs were cultured in EGM-2 media unless otherwise specified, consisting of 2% FBS, hEGF, hydrocortisone, VEGF, hFGF-B, R3-IGF-1, ascorbic acid, heparin, and gentamicin/amphotericin-B (Lonza, Walkersville, MD). MSCs were cultured in low glucose DMEM and L-glut media, with 10% FBS (Invitrogen). Cells were cultured in 5% CO_2_ at 37° C, and used at passage 5.

### 2.2. GelMA hydrogel formation

Gelatin (Sigma) was functionalized with methacrylamide groups as described previously.^57^ We adapted our previously published method to reduce the amount of methacrylic anhydride used and control the pH during the reaction. Methacrylic anhydride (MA) was added dropwise to a solution of gelatin and carbonate-bicarbonate (CB) buffer at a pH of 9-10.^63^ After an hour, the reaction was stopped by adding warm DI water. The pH was checked to ensure it was between 6-7, then the gelatin solution was dialyzed against DI water for one week and lyophilized. The gelatin:MA ratio was altered to control the degree of functionalization (DOF), and quantified by H^1^ NMR. For this study, GelMA with a DOF of 55% was used for all experiments. Hydrogels were made by dissolving GelMA at 5wt% in PBS at 65° C, with 0.1% w/v lithium acylphosphinate (LAP) as photoinitiator. The prepolymer solution was pipetted into circular Teflon molds with a diameter of 10 mm and thickness of 2 mm for mechanical testing, or 5 mm diameter and 1 mm thickness for cell culture experiments, then exposed to UV light (λ = 365 nm, 7.1 mW cm^−2^) for 30s.

### 2.3. GelMA hydrogel mechanical testing

Hydrogels were tested to determine the Young’s modulus in unconfined compression using an Instron 5943 mechanical tester (Instron, Norwood, MA). Hydrated (PBS) samples were compressed with a strain rate of 0.1 mm min^-1^. The Young’s modulus was determined as the slope of a linear fit applied to 15% of the stress vs strain data, with an offset of 2.5% using a custom MATLAB code.

### 2.4. Fluorescence recovery after photobleaching (FRAP)

The diffusivity of the hydrogel was determined using fluorescence recovery after photobleaching (FRAP) with a confocal microscope (Zeiss LSM710 Multiphoton Confocal Microscope, Germany) and two sizes of FITC-dextran probes, 40kDa and 250 kDa. Acellular hydrogels were incubated overnight in 200 ug mL^-1^ FITC-dextran in PBS. A 405 nm laser was used to photobleach a circular spot (∼30 – 40µm dia.), with recovery monitored (via 488 nm laser) for up to 100 sec (**Fig. 1B**). Recovery data was then analyzed using a MATLAB code described previously in *Jönsson et al* to determine small molecule diffusivity (representative fit: **Fig. 1C**).^38^

**Figure 1:**
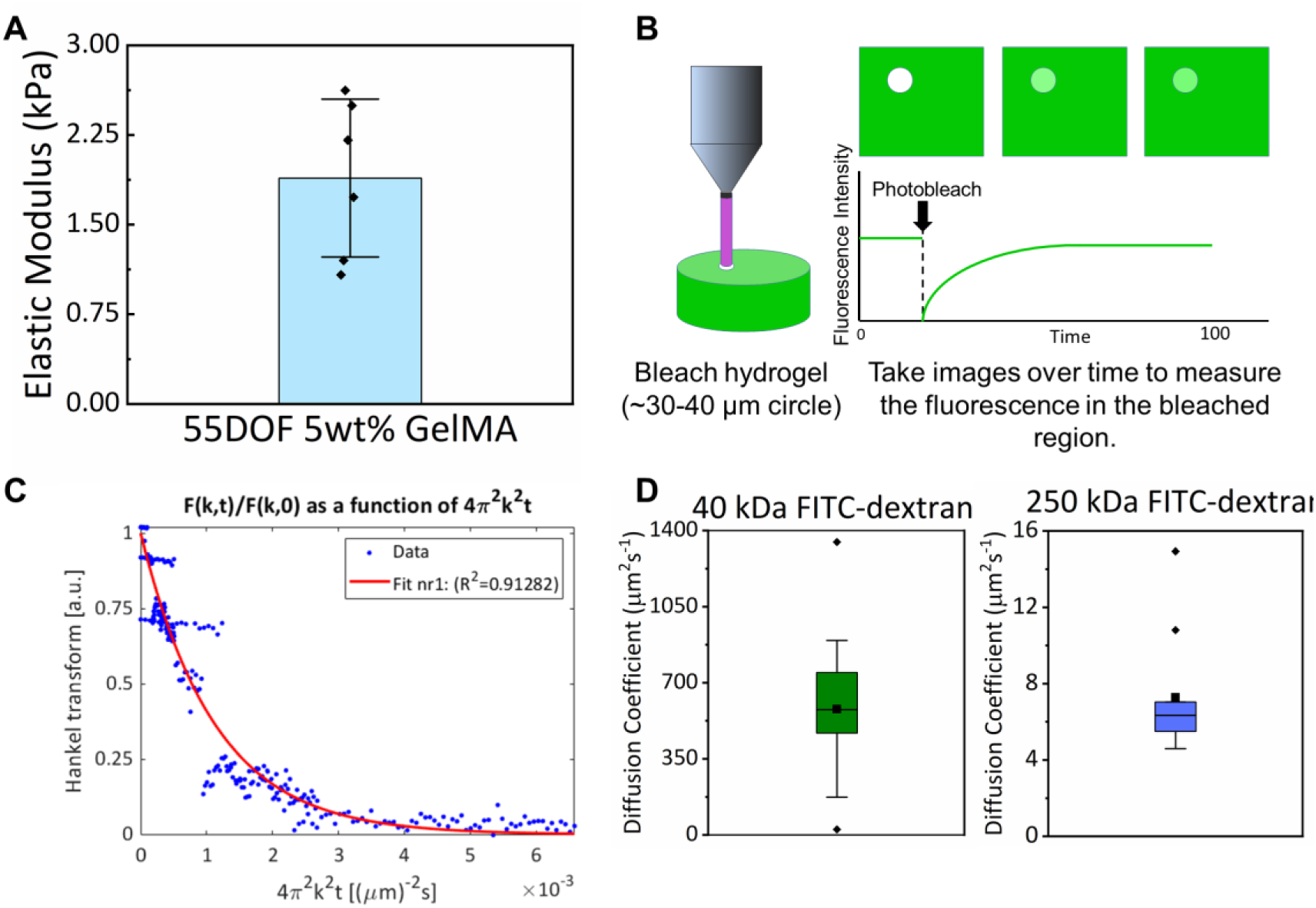
Physical properties of the GelMA hydrogel. A) Elastic modulus measured in compression for the gel formulation used in all experiments: 5wt%, 55 degree of functionalization GelMA. Data presented as Mean ± SD. B) Schematic of FRAP experiment. Hydrogels are soaked overnight in FITC-dextran, then locally bleached with a 405 nm laser, then time-lapse analysis of fluorescence recovery. C) Example of the raw data from the FRAP images (blue dots) and curve fit (red line) to calculate the diffusivity coefficient. D) Calculated diffusion coefficients for the FITC-dextran molecular weights of 40 kDa and 250 kDa. Data presented as a box plot.

### 2.5. Endothelial Network Formation

Endothelial networks were formed in hydrogels by co-culturing HUVECs and MSCs in a 2:1 ratio (HUVEC:MSC) with cell densities of either 1.5 × 10^6^ or 3 × 10^6^ total cells mL^-1^.^48^ Cells were suspended in prepolymer solution, pipetted into Teflon molds, and photopolymerized as described previously. Hydrogels were maintained for 7 days in EGM-2 media with daily media changes. On day 6, hydrogels were switched to endothelial basal media (EBM) with no serum or growth factors added. This allowed the collection of conditioned media without any added factors on day 7.

### 2.6. Immunofluorescent Staining of Endothelial Cell Networks

Hydrogels were fixed in formalin (Sigma Aldrich, St. Louis, MO) and washed with PBS. Gels were then soaked in PBS with 0.5% Tween 20 (Thermo Fisher Scientific, Waltham, MA) to permeabilize the encapsulated cells, and blocked in PBS with 2% bovine serum albumin (BSA) (Sigma Aldrich, St. Louis, MO) and 0.1% Tween 20. Endothelial cells were stained with mouse anti-human CD31 overnight at 4 °C, then the secondary of goat anti-mouse Alexa Fluor 488 overnight at 4 °C. PBS with 0.1% Tween was used to wash at all steps after permeabilization. Hoescht was added as a counterstain at 1:2000 in PBS for 30 min after the secondary.

### 2.7. Image Acquisition and Analysis

Networks were imaged with a DMi8 Yokogawa W1 spinning disk confocal microscope outfitted with a Hamamatsu EM-CCD digital camera (Leica Microsystems, Buffalo Grove, IL). Z-stacks were acquired with a depth of 200 µm and step size of 5 µm. Six regions were imaged per gel, three from each side of the hydrogel. The networks were quantified for metrics of total network length, and number of branches and junctions, using a previously established computational pipeline implemented in ImageJ and MATLAB.^16^ Briefly, a custom plugin in ImageJ filters and binarizes the z-stacks so that the vessels are extracted from the image. In MATLAB, the vessels are skeletonized and then converted into a quantifiable nodal network. As this approach generates a 3D skeleton of all vessels within an experimental volume, it is able to accurately resolve both the number of branches and vessel segments.^17^

### 2.8 Hematopoietic Stem and Progenitor Cell Isolation and Flow Cytometric Analysis

Primary murine HSPCs were obtained from the bone marrow of the femur and tibia of C57BL/6 female mice, age 4–8 weeks (The Jackson Laboratory). All work involving murine cell extraction was conducted under approved animal welfare protocols (Institutional Animal Care and Use Committee, University of Illinois at Urbana-Champaign). The bones were gently crushed and washed to extract the bone marrow and filtered through a 40 µm sterile filter with 5% FBS in PBS while on ice. ACK lysis buffer was added to remove blood cells and quenched with PBS/FBS. Initial enrichment for HSPC was performed with the EasySep Mouse Hematopoietic Progenitor Cell Enrichment Kit and Magnet (Stemcell Technologies, CA). Further isolation was performed to obtain a population of the Lin^−^ Sca-1^+^ c-kit^+^ (LSK) cells using a BD FACS Aria II flow cytometer (**Supp. Fig. 2**).^12,73^ The LSK population was collected into 25% FBS in PBS on ice and immediately used for experiments. LSK antibodies (eBioscience, CA) were as follows: APC-efluor780-conjugated c-kit (1:160), PE-conjugated Sca-1 (0.3:100), and Lin: FITC-conjugated CD5, B220, CD8a, CD11b (1:100), Gr-1 (1:400), and Ter-119 (1:200).^12,73^

### 2.9 Hematopoietic Stem and Progenitor Cell Conditioned Media Culture

For culture with perivascular conditioned media, LSKs were encapsulated in GelMA hydrogels as described previously,^26^ with 6000 LSK/gel. Hydrogels were maintained in 48 well plates for 1-4 days with daily media changes. Media was either a 50:50 blend of conditioned media collected from perivascular hydrogel cultures and Stemspan SFEM (Stemcell Technologies) with a final concentration of 100ng mL^-1^ SCF (Peprotech) and 0.1% penicillin/streptomycin (PS), or a control media of just Stemspan SFEM with SCF and PS.

On Day 1 and 4, cells were collected for analysis via fluorescence-assisted cytometry, using a BD LSR Fortessa (BD Biosciences, San Jose, CA). Hydrogels were dissolved using 100 units collagenase type IV (Worthington Biochemical Corp) on a shaker in the incubator 37°C for 20 min, with gentle mixing by pipetting. Degradation was quenched with 1 mL PBS + 5% FBS and centrifuged at 300 rcf x 10 minutes. The collected pellet was resuspended in PBS + 5% FBS and stained with a cocktail of antibodies in order to determine the populations present, using surface markers to differentiate long term (LT-HSCs: CD34^lo^ CD135^lo^ Lin^−^ Sca1^+^c-kit^+^) and short term (ST-HSCs: CD34^hi^ CD135^lo^ LSK) HSCs, and multipotent progenitors (MPPs: CD34^+^ CD135^+^ LSK).^68,73^ Following staining, the cells were fixed and permeabilized with Foxp3/Transcription Factor Staining Buffer Set (ThermoFisher), then stained with Ki67 for proliferation status, and DAPI (1mg/mL, ThermoFisher). Cells were resuspended in PBS + 5% FBS and analyzed via Fluorescence-Assisted Cytometry (FACs) using a BD LSR Fortessa (BD Biosciences, San Jose, CA). Fluorescence minus one (FMO) controls were created using lysed whole bone marrow for gating. DAPI was used to distinguish cells from debris, with a 5,000 DAPI threshold (Gating strategy: **Supp. Fig. 2**). Antibodies were supplied by eBioscience (San Diego, CA) as follows: PE-conjugated Ki-67 (0.3:100), eFluoro660-conjugated CD34 (5:100), PE-CY5-conjugated CD135 (5:100), APC-efluor780-conjugated c-kit (1:160), PE-CY7-conjugated Sca-1 (0.3:100), and Lin: FITC-conjugated CD5, B220, CD8a, CD11b (1:100), Gr-1 (1:400,), and Ter-119 (1:200).

### 2.10 Cytokine Analysis

Conditioned media was collected on Day 7 from endothelial networks with 3 × 10^6^ cells and no serum or growth factors for the final day of culture, as described above. A Proteome Profiler Human XL Cytokine Array (R&D Systems, Minneapolis, MN) was used to identify what cytokines are secreted by the endothelial network co-cultures. Each array uses 500 µL of conditioned media, and membranes were imaged in an autoradiography film cassette with an exposure time of 2 min. Images were then analyzed in ImageJ using the microarray profile and measuring the integrated density of the dots. Then, a signal was determined based on the percentage intensity compared to the positive control reference spots, with at least 5% considered to indicate the presence of a factor.

STRING analysis, or Search Tool for the Retrieval of Interacting Genes/Proteins, is a database and online resource for known and predicted protein-protein interactions.^67^ After determining the proteins expressed in the perivascular secretome, all eighteen were input into the website to see how they interact. The lines connecting nodes have different colors based on how the interaction was determined, such as from databases or experimentally determined.

### 2.11 Statistics

For experiments with two groups, a Student’s two sample t-test was performed in OriginPro (OriginLab, Northampton, MA). Normality of the data was determined using the Shapiro-Wilkes test, and homoscedasticity using Levene’s test. For groups that failed to meet the assumption of normality, a Mann Whitney test was used as a nonparametric alternative to the t-test. Experiments used n=6 hydrogel samples per group for both endothelial network characterization and HSC conditioned media experiments. FRAP data was tested for outliers using Grubb’s test with a significance value of 0.05 and displayed as a boxplot. Significance was set at p<0.05. Data are represented as mean ± standard error of the mean (SE).

## 3. Results

### 3.1. GelMA Hydrogel Properties

We determined biophysiclal properties of the hydrogels using unconfined compression and measurement of small molecule biotransport. The Young’s modulus was found to be 1.89 kPa ± 0.66 (**Fig. 1A**). The coefficient of diffusion was determined via fluorescence recovery after photobleaching (FRAP) (**Fig. 1B**) for two different molecular weight FITC-dextran probes, 40 kDa and 250 kDa, for the hydrogel (5wt% 55 degree of functionalization) with values of 578.24 ± 286 µm^2^ s^-1^ and 7.27 ± 3 µm^2^ s^-1^ respectively (**Fig. 1D**).

### 3.2. Endothelial Network Formation

We subsequently examined the influence of endothelial cell seeding conditions within the hydrogel on resultant endothelial network formation. Endothelial networks were formed by culturing human umbilical vein endothelial cells (HUVECs) with mesenchymal stem cells (MSCs) in a 2:1 ratio (HUVECs:MSCs). After seven days of culture, gels were fixed and stained for CD31, a cell adhesion molecule expressed in endothelial cells, to visualize network formation. Successful formation of networks occurred at two cell densities, 3×10^6^ cells mL^-1^ and 1.5×10^6^ cells mL^-1^ (**Fig. 2A**). We quantified total vessel length, number of branches, and number of vessels within the hydrogel as a function of cell seeding density (**Fig. 2B**). Increasing cell density to 3×10^6^ cells mL^-1^ led to significant increases in all parameters. As a result, this condition was selected for further characterization and experiments moving forward.

**Figure 2:**
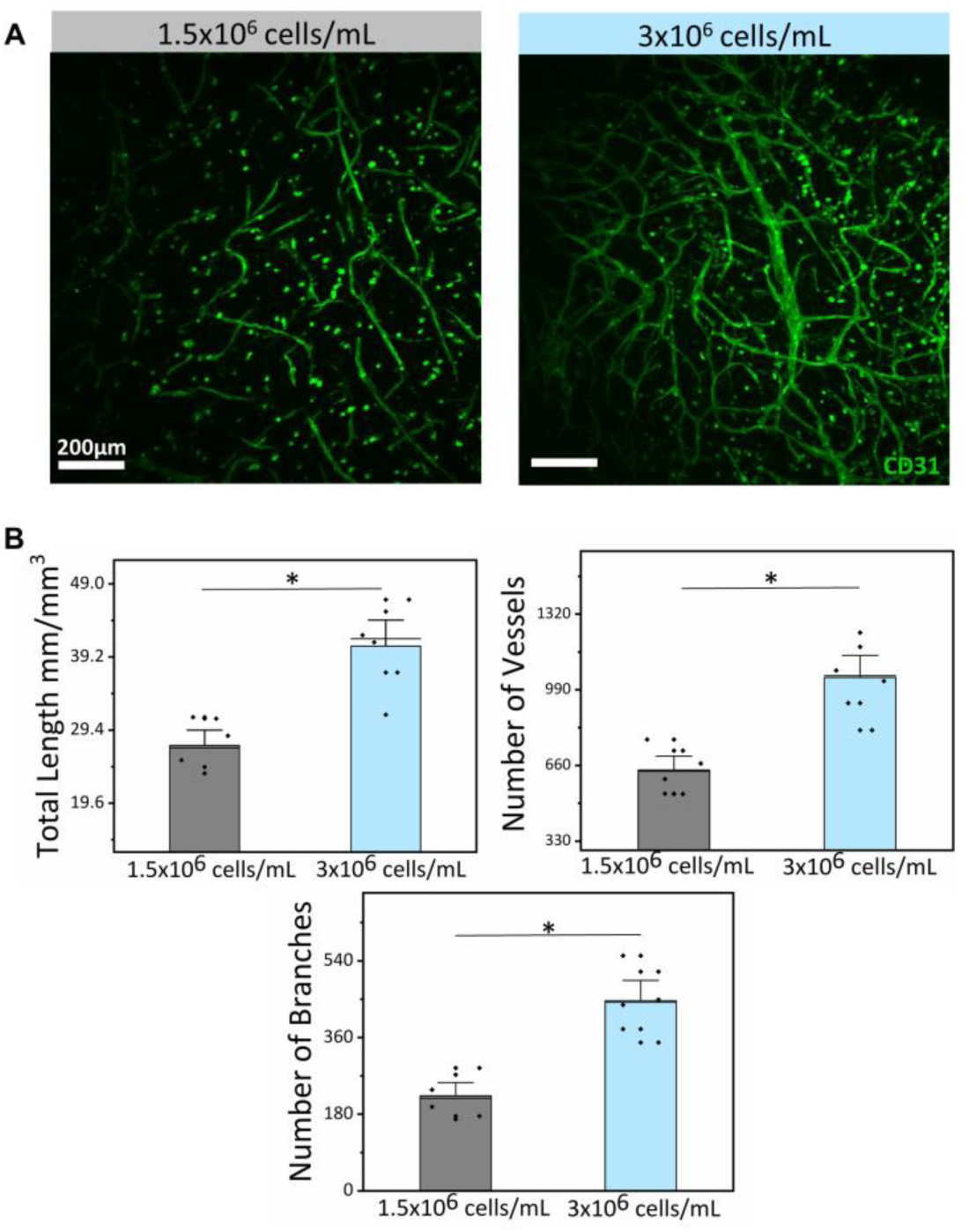
Engineered perivascular niche. A) Representative maximum intensity projections of CD31 staining of endothelial networks formed at cell densities of 1.5×10^6^ cells/mL or 3×10^6^ cells/mL (both 2:1 HUVEC:MSC). Scale bar: 200 μm. B) Network complexity metrics: total length, number of vessels, number of branches. Data represented as the mean ± SE. * represents p<0.05. n=6.

### 3.3. Secretome of the Perivascular Model

After successfully forming the networks, we characterized factors secreted by the perivascular cells during culture at day 7 of culture. To identify factors secreted by the endothelial networks, the conventional culture media was replaced at day 6 with media lacking all exogenous growth factors and serum, then isolated that media after 1 day of culture on the endothelial networks. We confirmed that serum removal from the media for that day did not affect the formation or stability of endothelial cell networks (**Supp. Fig. 1**). We subsequently performed a cytokine array that identified many cytokines expressed by the perivascular culture and that had also been identified via the literature as potentially important for HSC growth and/or maintenance (**Fig. 3A**). Several cytokines were detected via normalization to positive control spots, with a percentage of at least 5% indicating presence in the media (**Fig. 3B**). Many factors detected are suggested in the literature to influence HSC behavior, including Angiogenin (ANG),^28,64^ Angiopoietin-2 (ANG-2),^27,69^ Insulin-like growth factor binding protein-2 (IGFBP-2),^3,11^ Osteopontin (OPN),^59^ Serpin E1,^33,72^ vascular cell adhesion molecule-1 (VCAM-1),^45^ and dickkopf-1 (Dkk-1)^31^ (**Fig. 3C&D**). Several other factors were also found to be expressed by the perivascular cells, and while not necessarily found to be directly related to HSC function in the literature, were found to interact via STRING analysis (**Fig. 3C**). This could indicate possible indirect effects on cell types in the niche, not necessarily the HSCs themselves.

**Figure 3:**
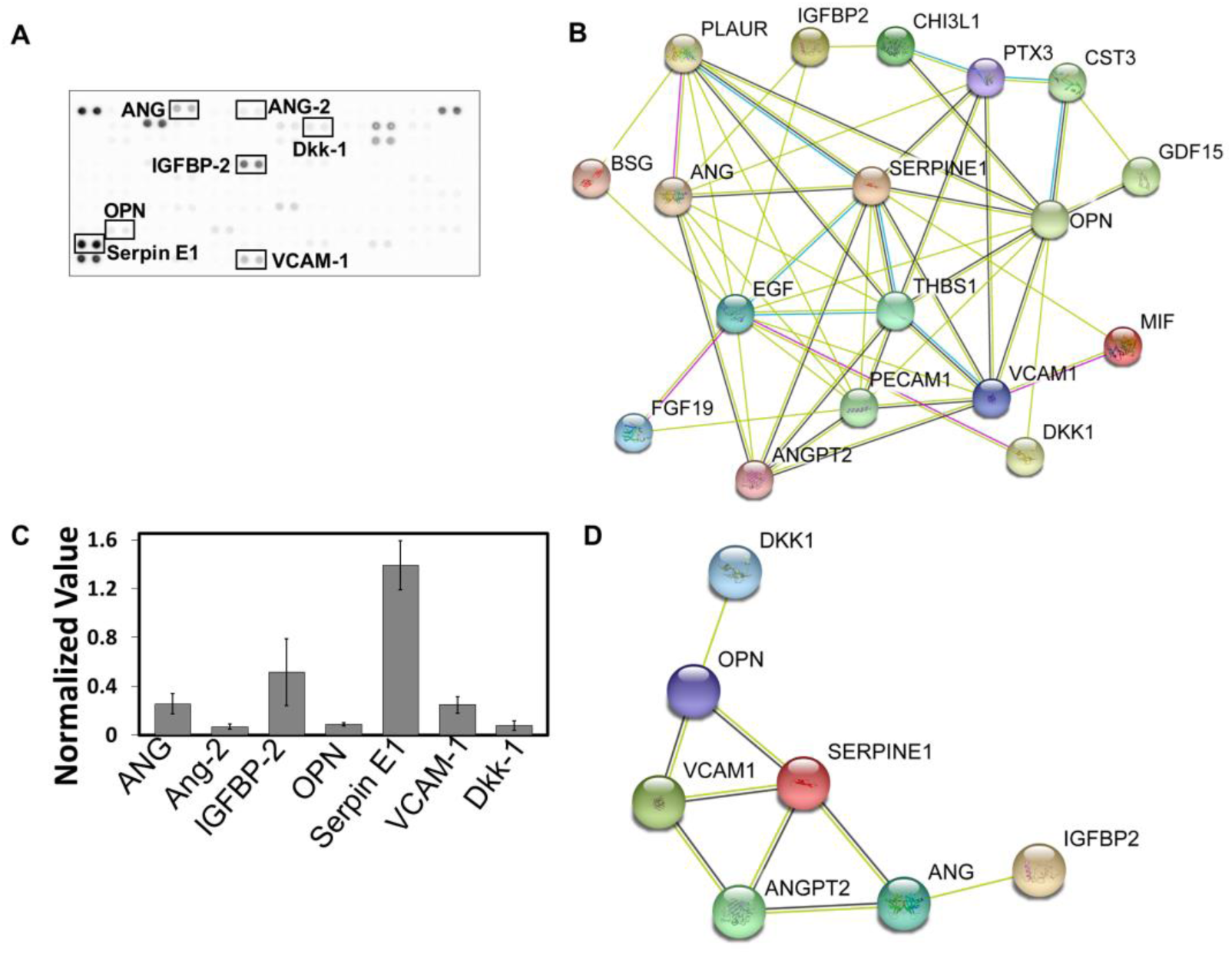
Analysis of the perivascular secretome. A) Representative cytokine array (factors in literature as important for HSCs highlighted). B) String analysis of the interaction of all cytokines detected. Blue lines represent known interactions from curated databases, and pink represents experimentally determined interactions. Light green lines represent interactions determined by textmining, and black represents the co-expression of the two proteins. C) Intensity of the cytokine spots normalized to the positive control of the cytokine array to show the normalized values and give a relative expression level. D) String analysis of only HSC related cytokines as determined by literature search.

### 3.4 Effect of Perivascular Conditioned Media on Hematopoietic Stem Cells

After analyzing the conditioned media of perivascular cultures and determining they secreted factors important for HSC fate decisions, we tested if these soluble factors were sufficient to maintain primitive HSCs. We cultured HSCs in a GelMA platform for up to 4 days with a blend of conditioned media and normal HSC growth media supplemented with SCF, or with normal HSC growth media alone (control). Following culture at days 1 and 4, HSCs were analyzed using flow cytometric analysis (**Supp. Fig. 2**). At Day 1, no differences are seen in the fractions of early progenitor cells (Lin^-^Sca-1^+^c-Kit^+^ or LSK) (**Fig. 4A**) normalized to LSK and lineage positive (Lin^+^), short-term HSCs (ST-HSCs) or multipotent progenitors (MPP) normalized to LSK (**Fig. 4B**). While we did not see an effect of perivascular conditioned media on the ST-HSC or MPP populations at day 4 (**Fig. 4C**), the presence of endothelial network condition media did influence the number of hematopoietic stem and progenitor cells. Normalizing the number of LSK cells to the total hematopoietic population (LSK + Lin^+^ cells), we observe a significant increase in the presence of endothelial conditioned media, suggesting an artificial perivascular network can secrete factors that help maintain hematopoietic progenitor cells in vitro (**Fig. 4C**).

**Figure 4:**
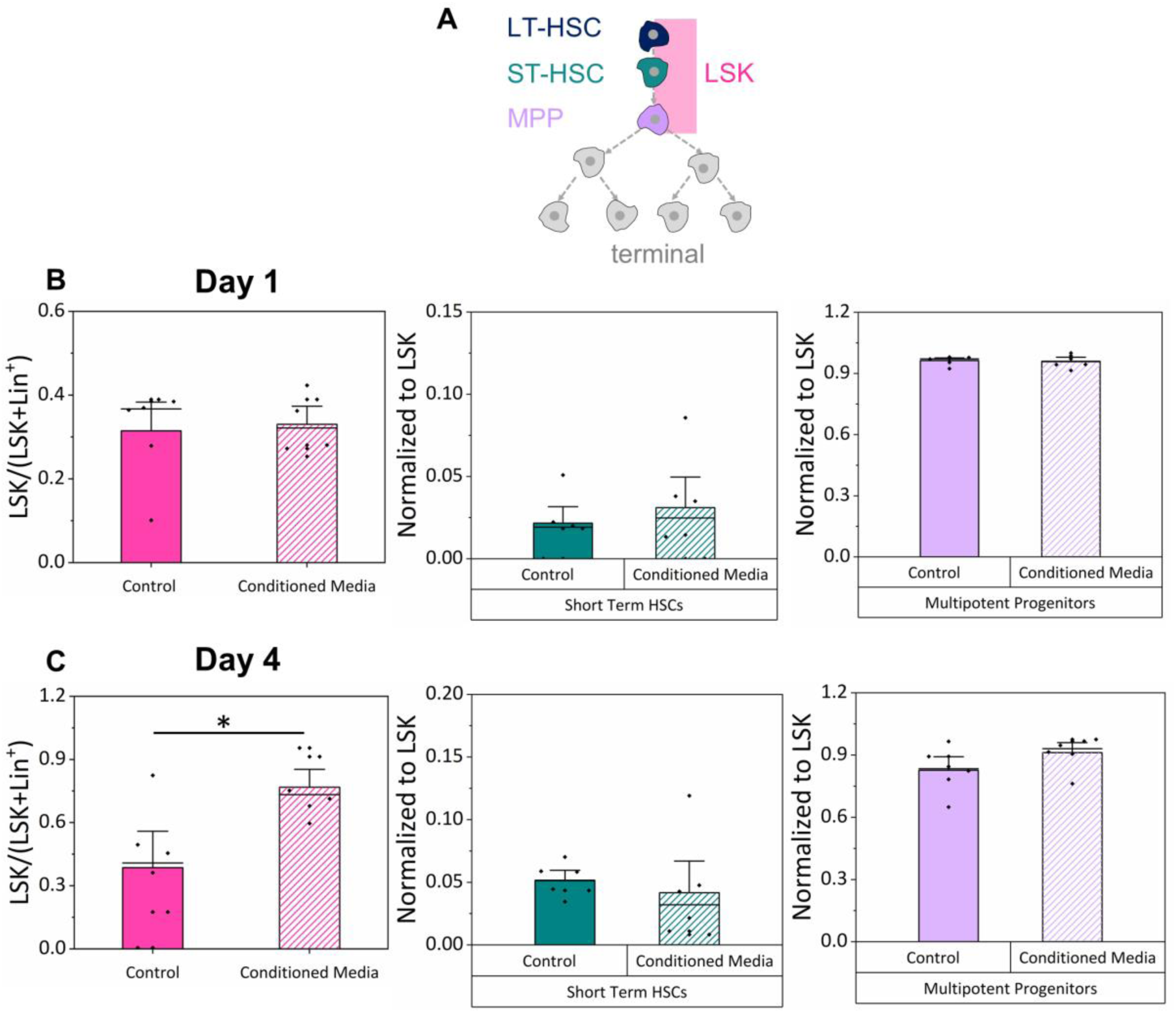
HSC culture with perivascular conditioned media. A) Lineage tree of HSCs. LSK population defined as the combination of LT-HSCs, ST-HSCs, and MPPs. B-C) Flow cytometry analysis of remaining cells after days 1 and 4. LSK numbers are normalized to the number of hematopoietic terminal cells (lineage positive Lin+) and LSKs combined. Short-term HSCs and multipotent progenitors are normalized to the number of LSKs. * p< 0.05.

## 4. Discussion

HSCs are a critical clinical tool due to their ability to reconstitute the entire blood and immune system. Improvements in the ability to culture and expand HSCs *in vitro* may lead directly to improved patient outcomes after transplant. Increasingly, a wide range of studies have attempted to move from identification of native niche features *in vivo*,^47,53,59,65^ to developing effective niche-mimicking conditioning procedures prior to transplant. Of these, the most popular have been addition of exogenous cytokines or culture with a feeder cell layer to expand HSC numbers, using factors shown *in vivo* to support HSC growth.^14,20,39,62,74^ In this work, we have shown the ability to form an artificial microvascular niche using endothelial networks that can secret factors important for HSCs. We further describe the role of isolated secretome on HSC culture as an essential first step for investigating the appropriate integration of cytokines into multi-dimensional biomaterial culture to facilitate HSC culture.

While the perivascular niche has emerged in recent studies as an important microenvironment for HSCs, many questions remain about how it supports HSCs.^2,8,53,61,71^ Technologies to examine multicellular cultures provide the potential to examine how HSCs relate to niche cells around them. Our lab has previously shown the use of biomaterial design principles to effectively regulate cellular crosstalk as a critical feature for *in vitro* HSC culture.^26,43^ In one investigation, we showed that while Lin+ niche cells isolated from the bone marrow generate a rich cytokine field to promote HSC myeloid differentiation and proliferation, adaptation of the hydrogel culture environment could significantly alter this response by altering the balance of paracrine signals from the Lin^+^ cells versus autocrine feedback generated endogenously by the HSC itself. Here, co-culture of HSCs with Lin^+^ cells in high diffusivity hydrogel environments that facilitate Lin^+^ to HSC paracrine signaling enhances HSC myeloid specification, while culture in low diffusivity hydrogels help maintain a primitive LSK population.^43^ More recently, we showed paracrine signals from marrow-derived mesenchymal stem cells could enhance the retention of quiescent HSCs,^26^ though this effect was also strongly dependent on the relative ratio of HSCs to MSCs as well as on processes of MSC-mediated hydrogel remodeling. Separately, a wide range of studies have focused on developing microphysiological models of the perivascular environment to interrogate processes associated with disease progression,^9,29,54,55^ cancer invasion or metastasis,^4,37,50,51,66^ and even dormancy of cancer stem cells.^10,24^ These studies provide an illustrative example of the power of the perivascular environment and motivate our study of the role of perivascular signals on murine HSC culture.

This work adapted technologies developed in our lab to create perivascular cultures using HUVECs and normal human lung fibroblasts (NHLFs) to study glioblastoma invasion and drug resistance.^48^ Here, we replaced traditional lung fibroblasts with human bone marrow derived MSCs, and report formation of robust endothelial networks that generate a rich secretome with multiple HSC related factors. We also adapted the generation of the methacrylamide-functionalized gelatin macromer we used to form GelMA hydrogels, reporting a GelMA hydrogel variant with 55% degree of functionalization as well as well-defined biophysical (modulus, diffusivity) properties. The GelMA hydrogel possesses marrow-associated moduli (native marrow: 0.1 to 10.9 kPa^36^) as well as well-defined diffusion properties. We used a well-established FRAP analysis approach^7,30,46,58^ using a range of dextran molecules at or larger than the size of the majority of cytokine molecular weights (40kDa). Our results suggest that while our hydrogel network is slowing small molecule diffusion (vs. water at room temperature: 2296 µm^2^s^-1^),^22^ diffusion in our GelMA hydrogel is largely consistent with diffusivities reported for other hydrogel systems.^30^ While altering parameters of the hydrogel such as weight percent or crosslinking could be used to adapt biotransport properties, they would also likely significantly impact the individual activity of both HSCs and the perivascular cultures.^26^ As a result, future studies will focus on co-optimization of GelMA hydrogel culture for individual HSC and perivascular culture metrics as well as for coordinated HSC-perivascular interactions. However, the important outcome from this work is that soluble HSC-perivascular interactions are an important consideration in biomaterial design criteria for generating artificial stem cell niche culture platforms.

This study only focused on the effect of secreted factors on HSC fate decisions and did not investigate direct HSC-perivascular interactions. Recent work by our group^49,50^ and *Ghajar et al*.^10,24^, in separate studies of the role of perivascular signals on glioblastoma invasion and drug resistance or breast cancer dormancy, have shown the importance of direct cell-cell contact as a critical axis of perivascular niche signaling. Such studies motivate ongoing efforts in our group to examine the role of indirect (secretome) versus direct cell-cell or matrisome mediated signaling. Opportunities also exist to interrogate HSC-perivascular interactions using primary cell types in co-culture to answer whether physical contact adds any additional support to HSC maintenance. Few *in vitro* studies have examined this question with primary, marrow derived perivascular niche cells in a physiologically relevant environment. *Poulos et al* co-cultured HSCs and vascular cells and found a benefit to co-culture over conditioned media, however this was performed in 2D cultures, lacking an instructive matrix environment.^60^ Similarly, *Kobayashi et al* cultured HSCs with primary endothelial cells and found that Akt activated endothelial cells maintained a greater HSC population than MapK activated endothelial cells, demonstrating the need for direct cell contact to support and expand of HSCs.^40^ However, this study utilized 2D cultures and had species mismatch in cell type. The mechanisms of these direct cell interactions remain poorly understood and represent an opportunity to use a hydrogel model system for further elucidation.

There are also opportunities to pursue a more targeted investigation of the secretome generated by the perivascular culture and its role on HSC activity. The secreted analysis performed here was qualitative rather than quantitative, thus provided no information about the concentration of different factors. However, it was essential to show that an engineered perivascular network could generate a broad spectrum of factors that are known to have an explicit role in hematopoietic stem cell activity. Many of the factors we identified in this study have been reported in the literature to effect HSC quiescence and cell activity. Angiogenin is suggested to play a role in the quiescence and self-renewal of HSCs, particularly LT-HSCs, and deletion of Ang from various niche cells led to more active cycling of LT-HSCs, ST-HSCs, and MPPs.^28,64^ IGFBP-2 has been studied as an exogenous factor *ex vivo* to expand HSCs, both on its own and in combination with SCF, thrombopoietin, or Flt-3 ligand.^11^ Additionally, IGFBP-2 has been suggested to support the survival and cycling of HSCs.^32^ Serpin E1 has conflicting roles in the HSC literature, with some finding it to be important for HSC retention in the niche and others finding its inhibition to aid hematopoietic regeneration;^33,72^ our identification of Serpin E1 suggests that a well-characterized biomaterial platform may be valuable to better elucidate the activity of factors with competing observations in vivo. VCAM-1 has been suggested to be important for HSC retention and expressed by perivascular MSCs and endothelial cells in the niche.^34,59^ Osteopontin is also expressed by perivascular MSCs and suggested to be important for the quiescence of HSCs, as well as their lodgment and engraftment into the niche.^52^ We also detected a factor that is suggested to have an inhibitory effect on HSCs: Dkk-1. Dkk-1 inhibits Wnt signaling, and inhibition of this pathway leads to increased cell cycling and a decline in regenerative function.^23^ Taken together, the connection between factors identified in our secretome screen and the available literature are consistent with our observation that perivascular conditioned media may help maintain a greater stem fraction of HSCs during *in vitro* culture. Future studies could use quantitative secretome screens paired with multidimensional analysis techniques such as Partial Least Squares Regression to reduce the dimensionality of complex data and help assign observational data to discrete biological processes.^35,41,70^ Indeed, we have recently shown that such approaches can help prioritize the study of individual factors from a secretome screen of marrow derived mesenchymal stem cells – identifying MSC-generated secretome may also help boost HSC quiescence.^25^ Future work integrating marrow derived MSCs and a perivascular niche may offer particular value for HSC expansion and quiescence.

The perivascular niche is recognized as an important regulator of HSC growth and function; however, there are still questions as to the important components of the microenvironment that lead to this effect. In this study, we show biomolecular signals generated by an engineered perivascular network can influence patterns of maintenance versus differentiation of primary murine hematopoietic stem and progenitor cells. We show the ability to monitor the formation of stable microvascular networks *in vitro*, analyze the content and hematopoietic-relevance of the secretome generated by these endothelial cell networks, then examine the role of these factors on the maintenance of primary HSCs *in vitro*. Notably, perivascular conditioned media promoted expansion of the hematopoietic stem and progenitor cells (LSK) population relative to the total number of hematopoietic cells in our 3D hydrogel model. Together, this suggests that biomaterial containing engineered perivascular niches may hold significant promise in the design of an engineered bone marrow niche for long term culture and expansion of HSCs.

## Acknowledgements

The authors would like to acknowledge Dr. Barbara Pilas of the Roy J. Carver Biotechnology Center (Flow Cytometry Facility, UIUC) for assistance with bone marrow cell isolation and flow cytometry. Research reported in this publication was supported by the National Institute of Diabetes and Digestive and Kidney Diseases of the National Institutes of Health under Award Numbers R01 DK099528 (B.A.C.H), as well as by the National Institute of Biomedical Imaging and Bioengineering of the National Institutes of Health under Award Numbers R21 EB018481 (B.A.C.H.). The content is solely the responsibility of the authors and does not necessarily represent the official views of the NIH. The authors are also grateful for additional funding provided by the Department of Chemical & Biomolecular Engineering and the Institute for Genomic Biology at the University of Illinois at Urbana-Champaign. The authors would like to acknowledge Zona Hrnjak and Aidan Gilchrist for the development of a custom Matlab code for rapid analysis of mechanical testing data.

