## Supplemental figures for "Perivascular Secretome Influences Hematopoietic Stem Cell Maintenance in a Gelatin Hydrogel"

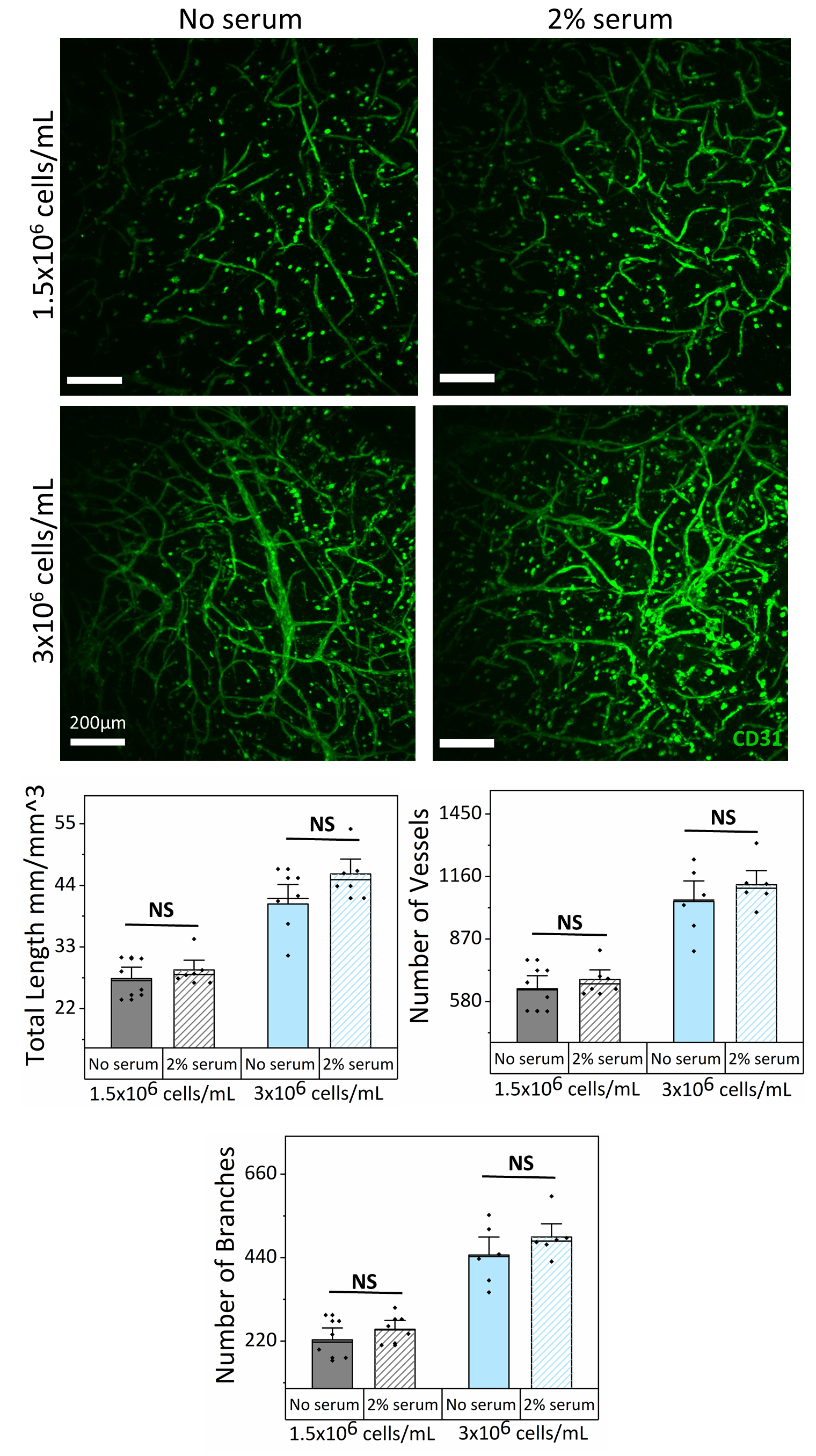

**Supplementary Figure 1.** Removing serum on Day 6 has no effect on network formation and stability. Maximum intensity projections of networks formed with cell densities of 1.5x10^6^ cells/mL and 3x10^6^ cells/mL with or without 2% serum for day 6-7 of culture. Network complexity measurements of total length, number of vessels and branches with and without 2% serum.

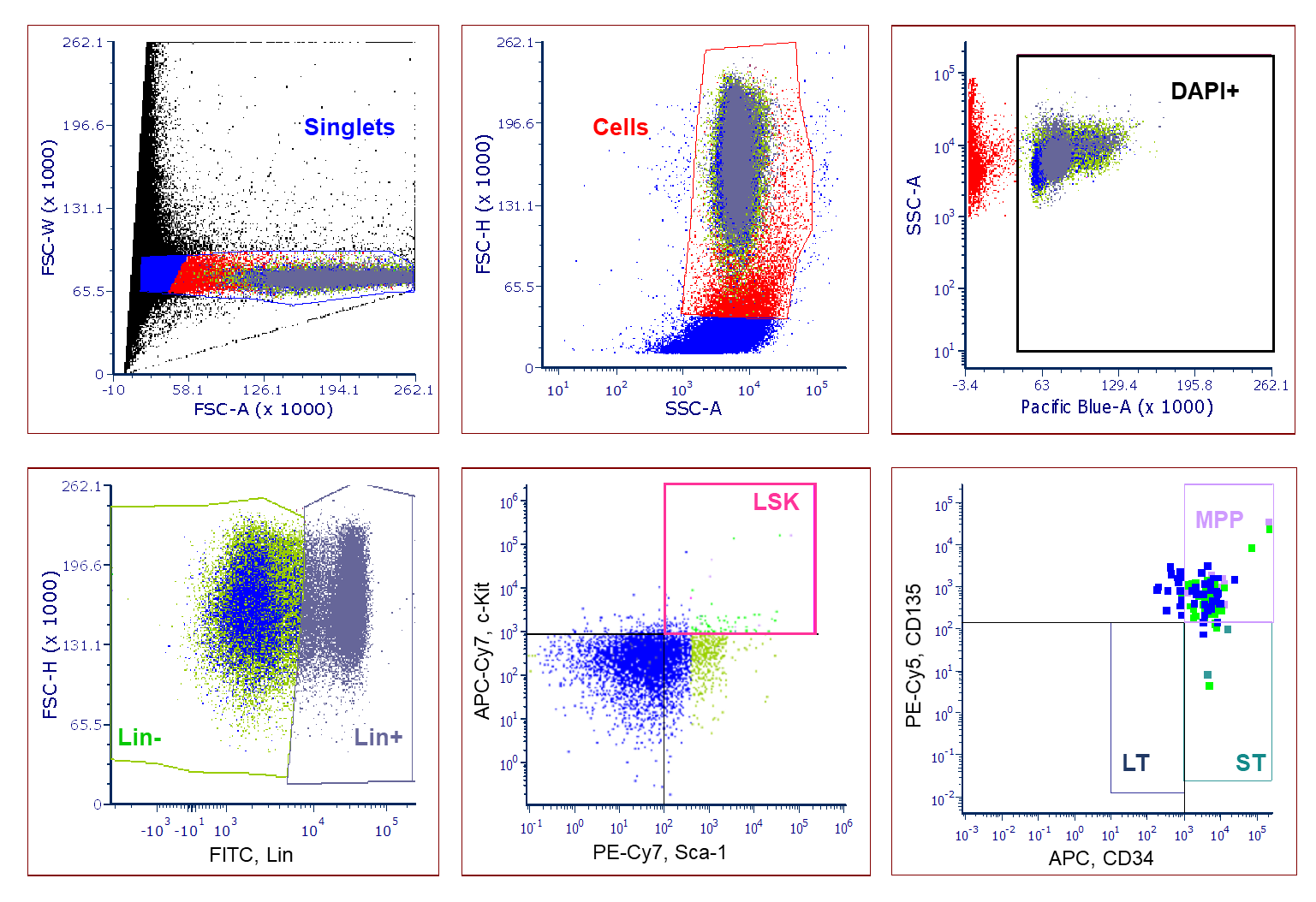

**Supplementary Figure 2.** Flow cytometry HSC gating strategy for data analysis after experiments. Data is first gated on singlets to remove particles that are likely not a single cell, then the data is gated by the size and shape of the particle to only include where cells should be found. Next, data is gated to only include DAPI positive particles to ensure the particle is a cell and not debris. Next, the data is gated into lineage negative (Lin-) and lineage positive (Lin+). The Lin- data is further gated for LSKs by taking the c-kit+ and Sca-1+ population, which can then be divided into the three groups of HSPCs, LT-, and ST- HSCs and MPPs based on their expression of CD34 and CD135.
